# Cumulative antibiograms: effects of isolate selection and stratification order on susceptibility estimates and resistance representation

**DOI:** 10.64898/2026.11.17.767230

**Authors:** Celso Soares Pereira Batista, Raúl Altaba, Daniela de Miniac, Dorotea Rodríguez, Ferran Navarro, Alba Rivera

## Abstract

Cumulative antibiograms depend on how isolates are selected and how data are stratified. Different selection strategies may affect both the number of isolates included and the susceptibility estimates reported. We retrospectively analyzed routine microbiology records from a tertiary-care hospital in Barcelona, Spain (2022-2025). Within each patient-organism-calendar-year unit, we compared three isolate-selection strategies: first-isolate, resistance-phenotype, and susceptibility-profile. We assessed the number of isolates retained, susceptibility at standard or increased exposure, the identification of resistance-phenotype profiles, and the effect of de-duplication before versus after stratification. Annual differences in susceptibility were estimated using paired cluster-bootstrap confidence intervals. For *Pseudomonas aeruginosa*, susceptibility-profile selection retained 16.9% more isolates than first-isolate selection, with differences exceeding 5 percentage points in 10 of 44 annual antimicrobial comparisons. No annual susceptibility differences exceeding this threshold were observed for the other species. Resistance-phenotype and susceptibility-profile selection criteria identified 468 MDR and 278 XDR *P. aeruginosa* isolates. Of these, 90 (19.2%) of the MDR and 53 (19.1%) of the XDR isolates were not captured by first-isolate selection. In the broader stratification dataset, stratification-first processing increased first-isolate retention by 21.2% for blood specimens and 11.2% for inpatients. These findings show that modest changes in cumulative susceptibility can coexist with incomplete representation of resistant phenotypes. In conclusion, phenotype- and AST-profile-based selection capture additional resistance information beyond first-isolate selection, despite modest differences in cumulative susceptibility. Clinical validation is needed before their use to guide empirical therapy.

**Importance:** Cumulative antibiograms summarize local susceptibility data to guide empirical antibiotic selection before individual susceptibility results are available. Including only the first isolate of each bacterial species per patient during a reporting period limits repeated counting but can leave other resistance patterns unrepresented. Our study showed that retaining additional isolates with different resistance patterns captured information absent from the first-isolate report, particularly for *Pseudomonas aeruginosa*. Small changes in overall susceptibility percentages did not necessarily reflect the extent of resistance information missed by first-isolate selection. Selecting isolates separately for blood samples or hospitalized patients also retained information lost when selection was performed across the entire dataset first. These findings support explicit reporting of selection rules and complementary analyses of resistance patterns. They do not show that additional isolates represent new infections or that changing the selection strategy improves treatment outcomes.

## Introduction

Cumulative antimicrobial susceptibility test (cAST) reports summarize routine antimicrobial susceptibility testing results to guide empirical antimicrobial selection, support antimicrobial stewardship, and monitor local antimicrobial resistance trends [1,2]. For routine antibiograms, the Clinical and Laboratory Standards Institute (CLSI) recommends including only the first isolate of each species per patient during the reporting period, regardless of specimen source or susceptibility profile [1]. This approach prevents repeatedly sampled patients from disproportionately influencing susceptibility estimates. The Spanish Antibiogram Committee (COESANT) similarly recommends first-isolate selection while recognizing the value of alternative analyses for specific resistance-related questions [3].

The impact of duplicate-isolate exclusion depends on the organism and selection criteria applied. Some studies have reported similar susceptibility estimates using patient- and episode-based approaches [4], whereas susceptibility-based criteria produced larger differences in others [5,6]. Time-based exclusion rules may have particularly marked effects for *P. aeruginosa* [7,8]. Differences between National Healthcare Safety Network (NHSN) and CLSI criteria have likewise shown that including additional isolates can increase reported non-susceptibility [9].

CLSI recognizes that first-isolate selection may omit subsequent resistance findings and describes supplemental analyses in which resistance profiles are counted once per patient [1]. However, changes in antimicrobial susceptibility percentages and the number of additional resistance-positive observations measure different features of a report.

Susceptibility percentages may remain similar even when additional resistant isolates or resistance profiles are identified. Conversely, including more isolates or resistance profiles does not necessarily provide a more accurate estimate of susceptibility. The results may still be influenced by how often patients are sampled, which antibiotics are tested, missing susceptibility results, and variability in antimicrobial susceptibility test (AST) measurements [1,10].

Stratification introduces a related issue; CLSI recommends applying first-isolate selection within the subset of interest [1]. Consequently, de-duplicating globally before stratification may yield a different population from stratifying first and then de-duplicating within each subgroup.

We compared three isolate-selection strategies using routine microbiology data from a tertiary-care hospital during 2022-2025: first-isolate selection, resistance-phenotype-based selection, and AST-profile-based selection. The study objectives were to quantify isolate retention under each strategy, assess differences in annual antimicrobial susceptibility estimates, and determine which resistance-positive observations were captured or missed and the timing of their detection. We also evaluated the impact of de-duplication and stratification order on isolate retention across patient settings, clinical departments, and specimen types.

## Materials and Methods

### Study design, data source, and eligibility

We conducted a retrospective laboratory-based methodological study at Hospital de la Santa Creu i Sant Pau, Barcelona, Spain, using routine microbiology records from January 2022 through December 2025. Data included patient, specimen, clinical-setting, organism, resistance phenotype, and categorical antimicrobial susceptibility testing (AST) variables. Diagnostic isolates were included; carriage-screening cultures were excluded. No formal sample-size calculation was performed.

### Microorganism identification and antimicrobial susceptibility testing

Organisms were identified by MALDI-TOF MS (Bruker Daltonik GmbH, Bremen, Germany). AST was performed by disk diffusion (I2A, France) and Sensititre broth microdilution (Thermo Fisher Scientific, United Kingdom) and interpreted using The European Committee on Antimicrobial Susceptibility Testing (EUCAST) clinical breakpoints applicable at the time of testing [11].

Extended-spectrum beta-lactamases (ESBL) production was assessed by double-disk synergy testing, AmpC β-lactamase by cloxacillin inhibition, following EUCAST guidance [12], and carbapenemase production by NG-Test CARBA 5 (NG Biotech, France), detecting KPC, VIM, IMP, NDM, and OXA-48-like enzymes.

Resistance phenotypes included ESBL, AmpC β-lactamase (plasmid-mediated and chromosomally de-repressed), and carbapenemase-producing Enterobacterales. For *P. aeruginosa*, multidrug-resistant (MDR) and extensively drug-resistant (XDR) definitions were adapted from the international criteria of Magiorakos *et al*. [13]. Methicillin-resistant *Staphylococcus aureus* (MRSA) and the penicillin-cefotaxime profile for streptococci were also included.

### Data preparation and susceptibility summaries

AST records were consolidated by calendar year, and the same dataset was used independently for all three selection strategies. Results were standardized as S, I, or R according to EUCAST [14]. S and I were combined as susceptible (S+I) for cumulative summaries and AST-profile comparisons, consistent with COESANT [3]. Missing AST results were excluded from antimicrobial-specific denominators.

For all three selection strategies and across all study years, inclusion was restricted to organisms or reporting groups with at least 30 isolates retained per year under the first-isolate strategy. This criterion ensured that the same organisms and reporting groups were evaluated across all strategies.

### Isolate-selection strategies

Selection was performed within patient-organism-calendar-year units, which were analytical units rather than clinically adjudicated infection episodes. The following selection strategies were evaluated:

#### First isolate (hereafter referred to as method X for brevity)

The earliest isolate per patient, organism, and year was retained regardless of specimen source or later susceptibility changes, using the AMR R package first-isolate algorithm [15], consistent with CLSI and COESANT [1,3].

#### Resistance phenotype (referred to as method Y)

The earliest representative of each distinct resistance phenotype profile was retained. Phenotypes were defined as ESBL, AmpC, and carbapenemase for Enterobacterales; MDR and XDR *P. aeruginosa*; MRSA; and penicillin-cefotaxime profile for streptococci.

#### AST profile (referred to as method Z)

The earliest representative of each distinct organism-specific AST profile was retained. For each antimicrobial, results were classified as resistant (R) or non-resistant (S+I) using antimicrobials available in both records; missing results alone did not define a new profile and were not included. Z was generated independently from the consolidated dataset, not from Y-selected isolates.

### Outcomes and timing analyses

We assessed differences in the number of isolates retained by methods Y and Z compared with method X, differences in cumulative susceptibility estimates, and resistance phenotypes not represented as positive by method X. We also examined the timing and characteristics of additional records and the effect of performing de-duplication before versus after stratification.

Resistance-positive observations were defined as unique patient-organism-calendar-year-phenotype combinations. For each resistance phenotype and comparator method (Y or Z), we calculated the proportion of positive observations not represented by method X. The denominator included all observations positive for that phenotype identified by the comparator, whereas the numerator included only those not represented as positive by method X. For these additional observations, time to detection was measured from the date of the X-selected isolate to the date of the first comparator-retained isolate exhibiting the corresponding resistance phenotype. Observations detected on the same day were assigned a time to detection of 0 days. Specimen-category concordance with X was summarized separately for observations occurring within and beyond 14 days. Concordance indicated that both observations belonged to the same specimen category, not that they originated from the same individual specimen.

### Order of de-duplication and clinical stratification

Two processing orders were compared for stratification by patient setting, clinical department, and specimen type. In order A, de-duplication was performed on the complete dataset before the retained isolates were assigned to their respective strata. In order B, isolates were first assigned to strata, and de-duplication was then performed independently within each stratum.

### Statistical analysis

Data were summarized as counts, percentages, and medians with inter-quartile ranges. Patient- and specimen-level summaries used unique patients and distinct specimens, respectively.

Retained-isolate counts were compared within patient-organism-year blocks using the Friedman test followed by paired Wilcoxon signed-rank tests with Holm adjustment. Absolute and relative gains were calculated versus X; pooled 95% confidence intervals (CIs) were estimated by patient-level bootstrap.

Cumulative susceptibility was S+I divided by all available S/I/R results for each antimicrobial. Annual paired differences were expressed in percentage points and accompanied by percentile 95% CIs from 1,000 cluster-bootstrap replicates using identical patient-organism-year resamples across strategies. Absolute differences >2, >5, and >10 percentage points were descriptive only; no P values were calculated for susceptibility comparisons.

Orders A and B were compared using two-sided exact binomial tests for paired discordant records and resistance-positive units. P values were adjusted by the Benjamini-Hochberg method within each strategy; adjusted P<0.05 was considered significant for retained-count and discordance comparisons.

### Software and reproducibility

Analyses were performed in R using AMR, dplyr, tidyr, rstatix, readxl, writexl, and openxlsx. The bootstrap seed was 20261004. Automated checks reconciled dates, retained-isolate counts, phenotype-positive units, and antimicrobial-specific denominators.

### Ethics statement

The study was approved by the Ethics Committee of Hospital de la Santa Creu i Sant Pau (IIBSP-WIS-2022-144; April 17, 2024) and conducted in accordance with the Declaration of Helsinki and European data protection regulations.

## Results

### Study population and specimen characteristics

The cohort comprised 18,674 unique patients across the four-year study period (2022-2025), contributing a total of 22,167 annual patient counts, as some patients were represented in multiple calendar years. Annual patient counts ranged from 5,314 to 5,875. Female patients accounted for 11,257 (60.3%) of the cohort, and 11,491(61.5%) patients were aged ≥65 years. During the study period, 10,590 patients (56.7%) contributed one specimen and 1,831 (9.8%) contributed at least five (Table 1).

**Table 1.** Patient and specimen characteristics.

| **a. Patient characteristics** |  |  |  |  |  |
| **Characteristic** | **Total** | **2022** | **2023** | **2024** | **2025** |
|---|---|---|---|---|---|
| **Patient characteristics** |  |  |  |  |  |
| Patients (unique per column) | 18,674 | 5,444 | 5,534 | 5,875 | 5,314 |
| Patient-years (sum of years) | 22,167 | 5,444 | 5,534 | 5,875 | 5,314 |
| **Age at first specimen, years** |  |  |  |  |  |
| <1 | 480 (2.6%) | 132 (2.4%) | 116 (2.1%) | 111 (1.9%) | 127 (2.4%) |
| 1-17 | 940 (5.0%) | 237 (4.4%) | 265 (4.8%) | 258 (4.4%) | 289 (5.4%) |
| 18-44 | 2,612 (14.0%) | 666 (12.2%) | 703 (12.7%) | 751 (12.8%) | 693 (13.0%) |
| 45-64 | 3,150 (16.9%) | 898 (16.5%) | 911 (16.5%) | 991 (16.9%) | 875 (16.5%) |
| 65-79 | 4,856 (26.0%) | 1,417 (26.0%) | 1,547 (28.0%) | 1,639 (27.9%) | 1,395 (26.3%) |
| ≥80 | 6,635 (35.5%) | 2,094 (38.5%) | 1,991 (36.0%) | 2,125 (36.2%) | 1,935 (36.4%) |
| Unknown | 1 (0.0%) | 0 (0.0%) | 1 (0.0%) | 0 (0.0%) | 0 (0.0%) |
| **Sex** |  |  |  |  |  |
| Female | 11,257 (60.3%) | 3,360 (61.7%) | 3,352 (60.6%) | 3,513 (59.8%) | 3,110 (58.5%) |
| Male | 7,415 (39.7%) | 2,084 (38.3%) | 2,181 (39.4%) | 2,361 (40.2%) | 2,204 (41.5%) |
| Unspecified | 2 (0.0%) | 0 (0.0%) | 1 (0.0%) | 1 (0.0%) | 0 (0.0%) |
| **Specimens per patient** |  |  |  |  |  |
| 1 | 10,590 (56.7%) | 3,531 (64.9%) | 3,539 (64.0%) | 3,652 (62.2%) | 3,387 (63.7%) |
| 2 | 3,583 (19.2%) | 1,001 (18.4%) | 1,010 (18.3%) | 1,077 (18.3%) | 904 (17.0%) |
| 3 | 1,695 (9.1%) | 417 (7.7%) | 420 (7.6%) | 487 (8.3%) | 465 (8.8%) |
| 4 | 975 (5.2%) | 198 (3.6%) | 215 (3.9%) | 219 (3.7%) | 225 (4.2%) |
| ≥5 | 1,831 (9.8%) | 297 (5.5%) | 350 (6.3%) | 440 (7.5%) | 333 (6.3%) |
| **b. Specimen characteristics** |  |  |  |  |  |
| **Characteristic** | **Total** | **2022** | **2023** | **2024** | **2025** |
| Distinct specimens | 41,288 (100.0%) | 9,703 (100.0%) | 10,112 (100.0%) | 11,455 (100.0%) | 10,018 (100.0%) |
| **Specimen category** |  |  |  |  |  |
| Blood | 7,311 (17.7%) | 1,759 (18.1%) | 1,856 (18.4%) | 2,016 (17.6%) | 1,680 (16.8%) |
| Urine | 17,079 (41.4%) | 4,521 (46.6%) | 4,213 (41.7%) | 4,480 (39.1%) | 3,865 (38.61lower %) |
| Lower respiratory | 4,174 (10.1%) | 843 (8.7%) | 1,024 (10.1%) | 1,217 (10.6%) | 1,090 (10.9%) |
| Cerebrospinal fluid | 90 (0.2%) | 21 (0.2%) | 32 (0.3%) | 19 (0.2%) | 18 (0.2%) |
| Other specimens | 12,634 (30.6%) | 2,559 (26.4%) | 2,987 (29.5%) | 3,723 (32.5%) | 3,365 (33.6%) |
| **Patient setting** |  |  |  |  |  |
| Inpatient | 16,305 (39.5%) | 3,630 (37.4%) | 4,005 (39.6%) | 4,617 (40.3%) | 4,053 (40.5%) |
| Outpatient | 24,983 (60.5%) | 6,073 (62.6%) | 6,107 (60.4%) | 6,838 (59.7%) | 5,965 (59.5%) |
| **Department** |  |  |  |  |  |
| Medical | 28,557 (69.2%) | 6,887 (71.0%) | 6,974 (69.0%) | 7,877 (68.8%) | 6,819 (68.1%) |
| Pediatrics | 2,152 (5.2%) | 524 (5.4%) | 534 (5.3%) | 538 (4.7%) | 556 (5.6%) |
| Surgery | 8,950 (21.7%) | 1,915 (19.7%) | 2,160 (21.4%) | 2,655 (23.2%) | 2,220 (22.2%) |
| Intensive care | 1,629 (3.9%) | 377 (3.9%) | 444 (4.4%) | 385 (3.4%) | 423 (4.2%) |

Patient-level percentages use unique patients as denominators; specimen-level percentages use distinct specimens. A patient may contribute to more than one annual column. Age was assigned at the first available specimen in the relevant overall or annual period. Specimen categories follow laboratory coding.

The dataset contained 41,288 distinct specimens, including 17,079 urine (41.4%), 7,311 blood (17.7%), 4,174 lower respiratory (10.1%), and 90 cerebrospinal fluid (0.2%). Outpatient and inpatient specimens accounted for 24,983 (60.5%) and 16,305 (39.5%), respectively; 1,629 specimens (3.9%) were assigned to intensive care (Table 1).

### Isolates retained by the three selection strategies

The 20 individually eligible organisms contributed 26,967, 27,632, and 29,294 isolates under methods X, Y, and Z, respectively. Relative to X, Y retained 665 more isolates (2.5%) and Z retained 2,327 more (8.6%). The 11 eligible Enterobacterales organisms contributed 17,756, 18,161, and 19,277 isolates, corresponding to gains of 405 (2.3%) with Y and 1,521 (8.6%) with Z (Table 2).

**Table 2.** Isolates retained by selection strategy.

| Organism | X, n | Y, n | Z, n | Y−X, n (%) | Z−X, n (%) |
|---|---|---|---|---|---|
| **Enterobacterales** |  |  |  |  |  |
| *Citrobacter freundii* | 137 | 139 | 140 | 2 (1.46%) | 3 (2.19%) |
| *Citrobacter koseri* | 281 | 281 | 284 | 0 (0.00%) | 3 (1.07%) |
| *Enterobacter cloacae* complex | 826 | 848 | 899 | 22 (2.66%) | 73 (8.84%) |
| *Escherichia coli* | 9,883 | 10,139 | 10,812 | 256 (2.59%) | 929 (9.40%) |
| *Klebsiella aerogenes* | 234 | 236 | 254 | 2 (0.85%) | 20 (8.55%) |
| *Klebsiella oxytoca* | 375 | 375 | 382 | 0 (0.00%) | 7 (1.87%) |
| *Klebsiella pneumoniae* | 3,293 | 3,399 | 3,673 | 106 (3.22%) | 380 (11.54%) |
| *Morganella morganii* | 426 | 427 | 434 | 1 (0.23%) | 8 (1.88%) |
| *Proteus mirabilis* | 1,814 | 1,830 | 1,896 | 16 (0.88%) | 82 (4.52%) |
| *Providencia stuartii* | 127 | 127 | 133 | 0 (0.00%) | 6 (4.72%) |
| *Serratia marcescens* | 360 | 360 | 370 | 0 (0.00%) | 10 (2.78%) |
| Total enterobacterales | 17,756 | 18,161 | 19,277 | 405 (2.3%) | 1,521 (8.6%) |
| **Pseudomonas** |  |  |  |  |  |
| *Pseudomonas aeruginosa* | 2,633 | 2,794 | 3,079 | 161 (6.11%) | 446 (16.94%) |
| **Staphylococci** |  |  |  |  |  |
| *Staphylococcus aureus* | 3,095 | 3,136 | 3,233 | 41 (1.32%) | 138 (4.46%) |
| *Staphylococcus epidermidis* | 1,305 | 1,360 | 1,496 | 55 (4.21%) | 191 (14.64%) |
| *Staphylococcus haemolyticus* | 227 | 227 | 231 | 0 (0.00%) | 4 (1.76%) |
| **Streptococci** |  |  |  |  |  |
| *Streptococcus agalactiae* | 449 | 449 | 450 | 0 (0.00%) | 1 (0.22%) |
| *Streptococcus anginosus* | 379 | 379 | 385 | 0 (0.00%) | 6 (1.58%) |
| *Streptococcus mitis* | 225 | 227 | 230 | 2 (0.89%) | 5 (2.22%) |
| *Streptococcus pneumoniae* | 593 | 594 | 604 | 1 (0.17%) | 11 (1.85%) |
| *Streptococcus pyogenes* | 305 | 305 | 309 | 0 (0.00%) | 4 (1.31%) |
| **Total** | **26,967** | **27,632** | **29,294** | **665 (2.5%)** | **2,327 (8.6%)** |

X, first-isolate selection; Y, resistance-phenotype selection; Z, AST-profile selection. Eligible organisms had at least 30 X-selected isolates in every study year. Counts are sums of annual selections, not unique patients over four years. Relative gains use X as the denominator. The 20 individually eligible organisms are included; overlapping cAST reporting groups are not summed.

For *P. aeruginosa*, X retained 2,633 isolates, Y retained 2,794, and Z retained 3,079. Relative gains were 161 isolates with Y (6.11%; 95% CI, 5.01-7.26%) and 446 with Z (16.94%; 95% CI, 14.82-19.23%). Among eligible staphylococcal species, Y retained 96 more isolates (2.1%) and Z retained 333 more (7.2%), including 191 additional *Staphylococcus epidermidis* isolates (14.6%) and 138 additional *Staphylococcus aureus* isolates (4.5%) with Z. Streptococcal gains were three isolates (0.2%) with Y and 27 (1.4%) with Z. *Escherichia coli* showed the largest absolute gain with Z (929 isolates; 9.4%), followed by *P. aeruginosa* and *Klebsiella pneumoniae* (380; 11.5%).

### Resistance-positive phenotypes not represented as positive by method X

Among Enterobacterales, Y identified 3,332 ESBL-positive, 666 AmpC-positive, and 94 carbapenemase-positive observations. Of these, 229 (6.9%), 39 (5.9%), and 12 (12.8%), respectively, were not represented as positive by method X. The corresponding values for method Z were 230/3,333 (6.9%), 41/668 (6.1%), and 12/94 (12.8%). For the additional positive observations identified by method Y, the median time to detection was 49.0 days (IQR, 18.0-117.0) for ESBL, 31.0 days (IQR, 6.0-54.0) for AmpC, and 48.5 days (IQR, 19.8-79.0) for carbapenemase production (Table 3).

**Table 3.** Resistance-positive analytical units not represented as positive by X.

| **Organism / phenotype** | **Y positive units** | **Y not positive under X, n (%)** | **Y interval, days, median (IQR)** | **Z positive units** | **Z not positive under X, n (%)** | **Z interval, days, median (IQR)** |
|---|---|---|---|---|---|---|
| Enterobacterales AmpC | 666 | 39 (5.9%) | 31.0 (6.0-54.0) | 668 | 41 (6.1%) | 25.0 (7.0-51.0) |
| Enterobacterales Carbapenemase | 94 | 12 (12.8%) | 48.5 (19.8-79.0) | 94 | 12 (12.8%) | 48.5 (19.8-79.0) |
| Enterobacterales ESBL | 3,332 | 229 (6.9%) | 49.0 (18.0-117.0) | 3,333 | 230 (6.9%) | 49.0 (18.2-117.0) |
| *P. aeruginosa* MDR | 468 | 90 (19.2%) | 32.5 (16.0-78.0) | 468 | 90 (19.2%) | 32.5 (16.0-78.0) |
| *P. aeruginosa* XDR | 278 | 53 (19.1%) | 42.0 (18.0-101.0) | 278 | 53 (19.1%) | 46.0 (22.0-101.0) |
| *S. aureus* MRSA | 840 | 25 (3.0%) | 76.0 (8.0-146.0) | 840 | 25 (3.0%) | 76.0 (8.0-146.0) |

Units are defined by patient–organism–calendar-year–phenotype and are not clinically adjudicated infection episodes. X, first-isolate selection; Y, resistance-phenotype selection; Z, AST-profile selection. Y- and Z-positive units are the method-specific denominators. “Not positive under X” denotes units not represented as positive by X. Intervals are measured from X to the first positive record retained by the comparator and are summarized only among units not represented as positive by X; day 0 is included. MDR includes XDR, and the two outcomes must not be summed. IQR, interquartile range; ESBL, extended-spectrum β-lactamase; MDR, multidrug resistant; XDR, extensively drug resistant; MRSA, methicillin-resistant *Staphylococcus aureus*.s

Both methods Y and Z identified 468 MDR-positive and 278 XDR-positive observations for *P. aeruginosa*. Of these, 90 (19.2%) and 53 (19.1%), respectively, were not represented as positive by method X. With method Y, the median time to detection of these additional resistance phenotypes was 32.5 days (IQR, 16.0-78.0) for MDR and 42.0 days (IQR, 18.0-101.0) for XDR. Six MDR-positive and three XDR-positive observations were detected on the same day as the X-selected isolate. With method Z, the median time to detection for XDR was 46.0 days (IQR, 22.0-101.0) (Table 3).

For *S. aureus*, both methods identified 840 MRSA-positive observations, of which 25 (3.0%) were not represented as positive by method X. The median time to detection was 76.0 days (IQR, 8.0-146.0), with five additional positive observations detected on the same day as the X-selected isolate (Table 3).

Of the 405 Enterobacterales records added by Y, 250 had at least one defined resistance phenotype when X had none, 114 had none when X was phenotype-positive, 34 were phenotype-positive at both assessments, and seven had no defined phenotype at either assessment. Among the 1,521 records added by Z, 656 had no defined phenotype at either assessment and 454 were phenotype-positive at both.

### Annual cumulative susceptibility estimates

Differences in annual cumulative susceptibility percentages (S+I, susceptible at standard or increased exposure) varied across organism groups. For Enterobacterales, 88 of 664 annual X-versus-Z comparisons (13.3%) showed absolute differences exceeding 2 percentage points, and 8 (1.2%) exceeded 5 percentage points. For *P. aeruginosa*, the corresponding figures were 29 of 44 (65.9%) and 10 of 44 (22.7%), with a median absolute difference of 3.46 percentage points. Among staphylococcal reporting groups, 10 of 332 comparisons (3.0%) exceeded 2 percentage points, but none exceeded 5 percentage points. No streptococcal comparison exceeded 2 percentage points (Table 4).

**Table 4.**
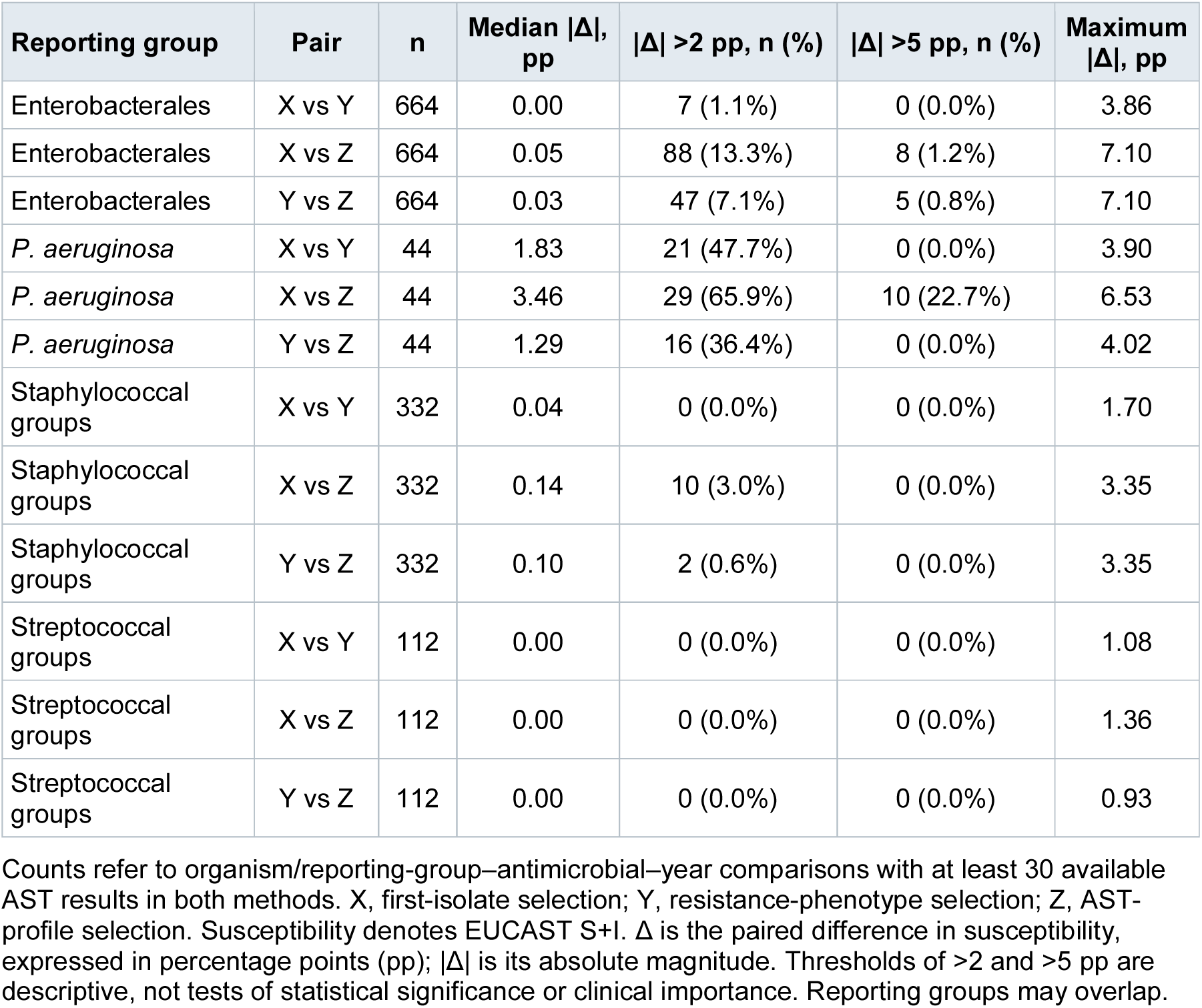
Annual susceptibility differences.

To illustrate the impact of isolate-selection strategies on cumulative susceptibility percentages, the 2024 *P. aeruginosa* antibiogram, which showed some of the largest differences, is presented in Table 5. Ciprofloxacin susceptibility was 78.1% with method X, 75.6% with Y, and 71.5% with Z, corresponding to a paired Z−X difference of −6.53 percentage points (95% CI, −8.42 to −4.74). For cefepime, the corresponding susceptibility percentages were 83.2%, 80.3%, and 77.8%, with a Z−X difference of −5.39 percentage points (95% CI, −7.62 to −3.45). Complete annual cumulative antibiograms are provided in Supplemental File S1.

**Table 5.** Cumulative antimicrobial susceptibility (S+I) test report of Pseudomonas aeruginosa, 2024.

| Organism | n | TZP* | CAZ* | FEP* | ATM* | IPM* | MEM | TOB | AMK | CIP* | COL | CZT |
|---|---|---|---|---|---|---|---|---|---|---|---|---|
| **Method X** |  |  |  |  |  |  |  |  |  |  |  |  |
| *P. aeruginosa* | 698 | **84.0%<br>(80.9-86.7)** | **85.4%<br>(82.7-88.0)** | **83.2%<br>(80.4-85.8)** | **95.4%<br>(93.8-96.8)** | **79.9%<br>(76.9-82.9)** | **96.7%<br>(95.3-98.0)** | **89.5%<br>(87.1-91.7)** | **98.6%<br>(97.6-99.4)** | **78.1%<br>(75.1-81.1)** | **99.9%<br>(99.6-100)** | **99.0%<br>(98.0-99.8)** |
| **Method Y** |  |  |  |  |  |  |  |  |  |  |  |  |
| *P. aeruginosa* | 741 | **81.4%<br>(78.3-84.0)** | **82.7%<br>(79.9-85.4)** | **80.3%<br>(77.4-82.9)** | **93.5%<br>(91.7-95.2)** | **77.8%<br>(74.6-81.1)** | **95.0%<br>(93.4-96.6)** | **88.7%<br>(86.3-91.0)** | **98.5%<br>(97.6-99.3)** | **75.6%<br>(72.4-78.7)** | **99.9%<br>(99.6-100)** | **98.6%<br>(97.5-99.5)** |
| **Method Z** |  |  |  |  |  |  |  |  |  |  |  |  |
| *P. aeruginosa* | 826 | **79.2%<br>(75.8-82.3)** | **80.3%<br>(76.9-83.2)** | **77.8%<br>(74.3-80.9)** | **92.6%<br>(90.2-94.7)** | **75.7%<br>(72.4-79.3)** | **93.1%<br>(91.1-95.1)** | **88.1%<br>(85.5-90.5)** | **98.3%<br>(97.2-99.2)** | **71.5%<br>(68.4-75.0)** | **99.6%<br>(99.2-100)** | **97.9%<br>(96.1-99.3)** |

Antibiotics: TZP, Piperacillin/tazobactam; CAZ, Ceftazidime; FEP, Cefepime; ATM, Aztreonam; IPM, Imipenem; MEM, Meropenem; TOB, Tobramycin; AMK, Amikacin; CIP, Ciprofloxacin; COL, Colistin; CZT, Ceftolozane/tazobactam.
Multidrug-resistant (MDR) and extensively drug-resistant (XDR) definitions were adapted from the international criteria of Magiorakos et al.
Method X: MDR: 112/698 (16.0%); XDR: 66/698 (9.5%).
Method Y: MDR: 145/741 (19.6%); XDR: 79/741 (10.7%).
Method Z: MDR: 177/826 (21.4%); XDR: 97/826 (11.7%).
Method X, first-isolate selection; Method Y, resistance-phenotype selection; Method Z, AST-profile selection
Microorganism-antibiotic combinations that require increased exposure are indicated with an asterisk (\*).
**<50%** **50-85%** **>85%**

### Timing and specimen context of additional records

To assess the timing of additional records retained by methods Y and Z, we compared their dates with those of the corresponding X-selected isolates. The analysis included 665 additional records retained by method Y and 2,327 by method Z. For method Y, 103 records (15.5%) were obtained on the same day as the X-selected isolate, 25 (3.8%) within 1-4 days, and 537 (80.8%) at least 5 days later. The corresponding figures for method Z were 415 (17.8%), 172 (7.4%), and 1,740 (74.8%), for the same time intervals (Table 6).

**Table 6.** Timing and specimen context of additional records.

| **Additional records** | **Y vs X** | **Z vs X** |
|---|---|---|
| Same-day (day 0) | 103 (15.5%) | 415 (17.8%) |
| At days 1–4 | 25 (3.8%) | 172 (7.4%) |
| After more than 4 days* | 537 (80.8%) | 1,740 (74.8%) |
| **Alternative** |  |  |
| Within 14 days | 186 (28.0%) | 793 (34.1%) |
| After 14 days | 479 (72.0%) | 1,534 (65.9%) |
| **Same specimen category** |  |  |
| Within 14 days | 147/186 (79.0%) | 559/793 (70.5%) |
| After 14 days | 359/479 (74.9%) | 1,213/1,534 (79.1%) |
| **Total** | **665 additional records** | **2,327 additional records** |

X, first-isolate selection; Y, resistance-phenotype selection; Z, AST-profile selection.
\*this number includes the records for “Within 14 days” and “After 14 days”
Counts in the first five rows are n (% of all additional records under the corresponding method). The ≤14-day and >14-day rows are an alternative partition of the same records; they must not be added to the day 0, day 1–4 and ≥5-day rows.
The final two rows are concordant/available comparisons (%), using the corresponding time-window denominator rather than all additional records. Time is measured from the X-selected isolate; day 0 is included. Concordance refers to specimen category, not to an identical specimen or a clinically adjudicated infection episode.

To explore whether additional records might reflect similar types of infection to those represented by the corresponding X-selected isolates, we compared their specimen categories as an indirect indicator of potential infection source. Comparisons were performed separately for records obtained within 14 days and more than 14 days after the X-selected isolate. Within 14 days, specimen categories matched in 147 of 186 comparisons (79.0%) for method Y and 559 of 793 (70.5%) for method Z. For records obtained more than 14 days later, the corresponding values were 359 of 479 (74.9%) and 1,213 of 1,534 (79.1%), for the same time intervals.

### Effect of stratification order

In the broader dataset used for the stratification analysis, de-duplicating within each stratum (order B), rather than before stratification (order A), increased the number of retained isolates across several clinical and specimen categories. For blood-specimens, order B retained 601 additional isolates with method X (21.2% increase), 570 (19.6%) with Y, and 477 (15.3%) with Z. For cerebrospinal fluid, 10 additional isolates were retained under each method, corresponding to increases of 21.3%, 20.8%, and 20.4%, respectively, although baseline counts were small. Among inpatients, the corresponding increases were 972 isolates (11.2%) with X, 912 (10.1%) with Y, and 778 (8.0%) with Z. In intensive care, the increases were 103 (12.0%), 105 (12.0%), and 90 (9.5%), respectively, whereas outpatient increases were 2.3%, 2.0%, and 1.5% (Table 7).

**Table 7.**
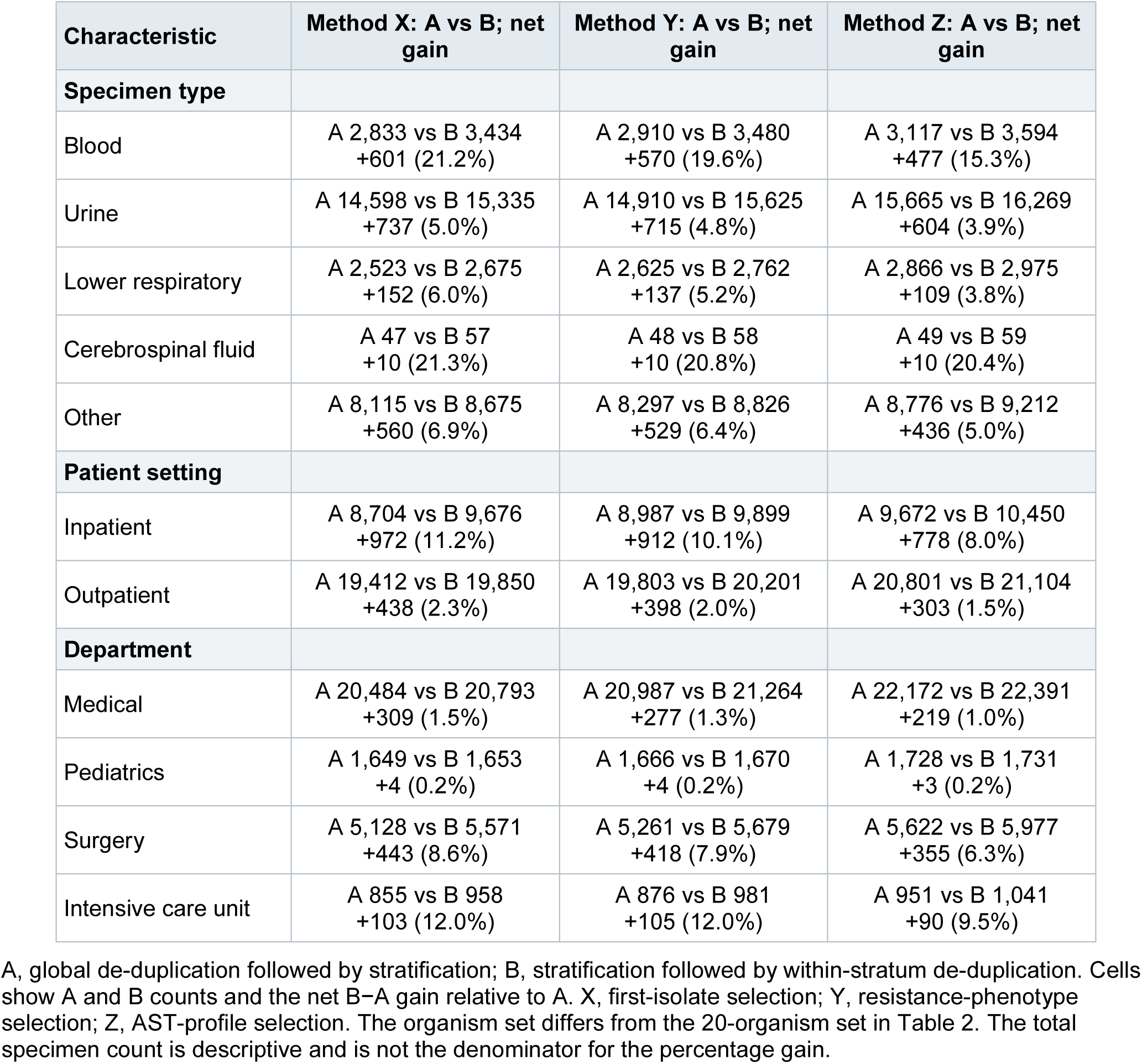
Records retained by stratification order.

| Characteristic | Method X: A vs B; net gain | Method Y: A vs B; net gain | Method Z: A vs B; net gain |
|---|---|---|---|
| **Specimen type** |  |  |  |
| Blood | A 2,833 vs B 3,434<br>+601 (21.2%) | A 2,910 vs B 3,480<br>+570 (19.6%) | A 3,117 vs B 3,594<br>+477 (15.3%) |
| Urine | A 14,598 vs B 15,335<br>+737 (5.0%) | A 14,910 vs B 15,625<br>+715 (4.8%) | A 15,665 vs B 16,269<br>+604 (3.9%) |
| Lower respiratory | A 2,523 vs B 2,675<br>+152 (6.0%) | A 2,625 vs B 2,762<br>+137 (5.2%) | A 2,866 vs B 2,975<br>+109 (3.8%) |
| Cerebrospinal fluid | A 47 vs B 57<br>+10 (21.3%) | A 48 vs B 58<br>+10 (20.8%) | A 49 vs B 59<br>+10 (20.4%) |
| Other | A 8,115 vs B 8,675<br>+560 (6.9%) | A 8,297 vs B 8,826<br>+529 (6.4%) | A 8,776 vs B 9,212<br>+436 (5.0%) |
| **Patient setting** |  |  |  |
| Inpatient | A 8,704 vs B 9,676<br>+972 (11.2%) | A 8,987 vs B 9,899<br>+912 (10.1%) | A 9,672 vs B 10,450<br>+778 (8.0%) |
| Outpatient | A 19,412 vs B 19,850<br>+438 (2.3%) | A 19,803 vs B 20,201<br>+398 (2.0%) | A 20,801 vs B 21,104<br>+303 (1.5%) |
| **Department** |  |  |  |
| Medical | A 20,484 vs B 20,793<br>+309 (1.5%) | A 20,987 vs B 21,264<br>+277 (1.3%) | A 22,172 vs B 22,391<br>+219 (1.0%) |
| Pediatrics | A 1,649 vs B 1,653<br>+4 (0.2%) | A 1,666 vs B 1,670<br>+4 (0.2%) | A 1,728 vs B 1,731<br>+3 (0.2%) |
| Surgery | A 5,128 vs B 5,571<br>+443 (8.6%) | A 5,261 vs B 5,679<br>+418 (7.9%) | A 5,622 vs B 5,977<br>+355 (6.3%) |
| Intensive care unit | A 855 vs B 958<br>+103 (12.0%) | A 876 vs B 981<br>+105 (12.0%) | A 951 vs B 1,041<br>+90 (9.5%) |

A, global de-duplication followed by stratification; B, stratification followed by within-stratum de-duplication. Cells show A and B counts and the net B−A gain relative to A. X, first-isolate selection; Y, resistance-phenotype selection; Z, AST-profile selection. The organism set differs from the 20-organism set in Table 2. The total specimen count is descriptive and is not the denominator for the percentage gain.

Stratification before de-duplication also increased the representation of resistance-positive observations. For blood specimens, resistance-phenotype-based selection and AST-profile-based selection under order B represented 135/493 (27.4%) and 91/493 (18.5%) ESBL-positive observations, respectively, that were not represented as positive under the corresponding order A analyses. Among inpatients, the corresponding figures were 131/902 (14.5%) and 82/902 (9.1%). For inpatient MRSA, the two selection methods under order B additionally represented 64/354 (18.1%) and 58/354 (16.4%) positive observations compared with order A. The corresponding values for MDR-positive *P. aeruginosa* were 22/284 (7.7%) and 15/284 (5.3%). Detailed results and stratified cumulative antibiograms for both processing orders are provided in the Table 8.

**Table 8.** Selected resistance-positive observations recovered by stratification-first processing.

| Stratum | Organism / phenotype | X: recovered / B-positive (%) | Y: recovered / B-positive (%) | Z: recovered / B-positive (%) |
|---|---|---|---|---|
| Blood | Enterobacterales / ESBL | 145/477 (30.4%) | 135/493 (27.4%) | 91/493 (18.5%) |
| Inpatient | Enterobacterales / ESBL | 159/863 (18.4%) | 131/902 (14.5%) | 82/902 (9.1%) |
| Inpatient | S. aureus / MRSA | 70/347 (20.2%) | 64/354 (18.1%) | 58/354 (16.4%) |
| Inpatient | P. aeruginosa / MDR | 35/233 (15.0%) | 22/284 (7.7%) | 15/284 (5.3%) |

A, selection on the complete dataset followed by stratification; B, stratification followed by selection within each stratum. X, first-isolate; Y, resistance-phenotype; Z, AST-profile selection.
Cells are B-positive patient–organism–calendar-year–phenotype observations not represented as positive by the corresponding A analysis, divided by all B-positive observations for that method and stratum. They are not percentages of all retained isolates or net isolate-count gains. MDR includes XDR; these categories must not be summed.

## Discussion

Isolate selection is not a neutral preprocessing step, because each strategy addresses a different clinical or epidemiological question. Method X, the first-isolate approach used for conventional cumulative antibiograms, provides standardized susceptibility estimates based on the first isolate per patient, organism, and year [1,3]. Method Y retains additional isolates when a predefined resistance phenotype is represented, asking whether clinically relevant resistance emerges later despite being absent from the X-selected isolate. Method Z retains distinct AST profiles, asking how much within-patient susceptibility-pattern diversity appears over time. In our study, changing the selection rule changed not only the number of retained isolates but also the representation of resistance phenotypes and the timing of their first detection. This is biologically plausible, because serial isolates may change susceptibility within days and the timing varies by organism and specimen source [16].

Compared with first-isolate selection, resistance-phenotype-based and AST-profile-based selection retained 2.5% and 8.6% more isolates, respectively. The largest increase was observed for *Pseudomonas aeruginosa*, for which AST-profile-based selection retained 16.9% more isolates than method X. These findings indicate that the impact of isolate-selection rules varies across organism groups and may be particularly pronounced for species with a greater potential for antimicrobial resistance development, such as *P. aeruginosa*. This organism has a well-recognized capacity to develop resistance through multiple intrinsic and acquired mechanisms, as well as adaptive changes in antimicrobial susceptibility [17]. Consequently, first-isolate selection may fail to capture part of the susceptibility-pattern diversity observed in this species.

Importantly, changes in cumulative susceptibility percentages did not necessarily reflect the extent of resistance information omitted by first-isolate selection. Although antimicrobial-specific susceptibility percentages (S+I) summarize the proportion of isolates categorized as susceptible at standard or increased exposure, they do not fully capture the occurrence of resistance phenotypes such as ESBL, carbapenemase production, MDR, or XDR. Consequently, clinically relevant resistance phenotypes may remain unrepresented in a cumulative antibiogram despite relatively small differences in overall susceptibility percentages. This distinction highlights the importance of evaluating not only changes in antimicrobial-specific susceptibility estimates but also the additional resistance information captured by alternative selection strategies.

The first-isolate rule limits the influence of repeatedly sampled patients and provides a standardized basis for comparisons across periods and institutions [1,3]. However, our findings also showed that additional isolates were not consistently more resistant than the initial isolate, indicating that later sampling does not necessarily yield a more resistant susceptibility profile. Discordant profiles among isolates obtained on the same day further illustrate the importance of selection criteria, as prioritizing the most resistant isolate, the most invasive specimen, or the earliest reported isolate may produce different estimates. Thus, alternative selection strategies should not be regarded simply as methods for identifying more resistant isolates, but rather as approaches that capture different dimensions of microbiological diversity.

Nevertheless, neither alternative strategy identifies distinct clinical infection episodes. Multiple resistance phenotypes or AST profiles may occur during a persistent infection, whereas a subsequent infection may remain unrepresented if its phenotype or AST profile has already been selected within the same patient-organism-year combination. The intervals between the X-selected isolates and additional records, which frequently extended over weeks or months, suggest that some additional isolates may correspond to separate clinical episodes. However, persistence, recurrence, colonization, strain replacement, and analytical variability cannot be distinguished without clinical information or strain typing. Therefore, a change in AST profile should not be interpreted as definitive evidence of resistance acquisition or a new infection [10].

The order of de-duplication and stratification affected both isolate retention and resistance-phenotype representation. In our study, stratification before de-duplication increased first-isolate retention by 21.2% for blood specimens and 11.2% for inpatients compared with de-duplication before stratification. These findings support the CLSI recommendation to perform stratification before removing duplicate isolates when constructing stratum-specific antibiograms [1]. This may be particularly relevant for setting- or syndrome-oriented reports, although specimen type and requesting service do not themselves define an infectious syndrome [18].

The implications of these findings are particularly relevant in the context of the recent debate on the role and future of hospital antibiograms. In their 2026 Point-Counterpoint article, Weiss and Basu questioned whether conventional annual antibiograms adequately address contemporary clinical and epidemiological needs [19]. The authors discussed limitations including the retrospective and aggregated nature of these reports, the potential under-representation of relevant resistance patterns, the influence of specimen source and patient population, and the limited capacity of hospital-wide susceptibility estimates to support individualized empirical treatment decisions. In contrast, they also emphasized that conventional antibiograms remain important for empirical treatment, antimicrobial stewardship, institutional surveillance, and formulary decisions, particularly when microbiological results are unavailable. An important distinction raised in this debate is that antibiograms intended to guide empirical therapy and analyses intended to monitor antimicrobial resistance do not necessarily require the same isolate-selection strategy [1–3,19]. CLSI M39 recommends standardized first-isolate selection for cumulative antibiograms, primarily to avoid over-representation of repeatedly sampled patients and facilitate comparisons across institutions and periods [1]. However, this approach was not designed to capture all within-patient changes in susceptibility or to provide comprehensive surveillance of resistance phenotypes. Alternative inclusion strategies may therefore be appropriate when the objective is to characterize resistance patterns, detect emerging phenotypes, or investigate changes in susceptibility over time. Our findings support this distinction, showing that first-isolate selection and phenotype- or AST-profile-based approaches provide complementary rather than interchangeable information.

Free, open-source tools such as AMR, R, and Python enable automated and reproducible antibiogram generation [15]. Development environments such as Positron and AI-assisted coding may facilitate their use by laboratories with limited programming expertise. However, automated analyses require validation to ensure methodological accuracy. Digital platforms could also facilitate access to customized antibiograms according to clinical setting and reporting objectives [19,20]. Nevertheless, their effect on prescribing decisions and patient outcomes requires prospective evaluation.

Transparent reporting of methodological decisions is essential for reproducibility and interpretation. Cumulative antibiograms should specify isolate-selection criteria, resistance- phenotype or AST-profile definitions, stratification order, same-day selection rules, and antimicrobial-specific denominators. Previous studies have identified incomplete reporting of these details [21]. Under EUCAST criteria, S+I estimates should be clearly defined, as category I indicates susceptibility at increased exposure [3,14]. Furthermore, changes in clinical breakpoints over time should be considered when comparing susceptibility trends, ideally by applying a consistent interpretative framework to historical data.

Future approaches should prioritize the integration of clinical data with antimicrobial susceptibility results to improve the clinical relevance of cumulative antibiograms, rather than relying solely on the methodological simplicity of selecting the first isolate per patient. Without such integration and clinical validation, the usefulness of conventional cumulative antibiograms for guiding empirical therapy may remain limited [19]. Ultimately, empirical treatment is selected for a clinical syndrome rather than an organism in isolation. Integrating microbiological data with infection source, severity, previous resistance, and other clinical variables would allow direct evaluation of episode- and syndrome-specific reports. Weighted- incidence syndromic combination antibiograms (WISCAs), which combine syndrome-specific organism frequencies with antimicrobial susceptibility data, may provide more direct estimates of empirical-regimen coverage than conventional organism-based reports [22].

This study has several limitations. It was conducted at a single center using routine categorical microbiology data without clinical adjudication of infection episodes, strain typing, or treatment-outcome assessment. Consequently, additional isolates could not be reliably classified as representing persistent infection, recurrence, or a new infection. The yield of phenotype- and AST-profile-based selection may depend on local sampling practices and testing panels, which may limit generalizability to other settings. Furthermore, the study assessed the microbiological information captured by different selection strategies but did not evaluate their predictive accuracy for empirical treatment or their impact on clinical outcomes.

In conclusion, modest differences in cumulative susceptibility can coexist with substantial amounts of additional resistance information not represented by first-isolate selection. Resistance-phenotype-based and AST-profile-based strategies provide a more comprehensive representation of local antimicrobial resistance by capturing resistance phenotypes and susceptibility profiles that would otherwise be overlooked. Validation against clinically defined episodes and treatment decisions is required before phenotype- or AST- profile-based selection can be considered a more accurate guide to empirical therapy.

## Data availability

Aggregate results and detailed cAST tables accompany the manuscript. Access to underlying pseudonymized routine records is available from the corresponding author on reasonable request, subject to applicable institutional and data-protection requirements. Sample implementations in R and Python is available on GitHub: https://github.com/drcelsobat/amr_selection_criteria

## Acknowledgments Funding

No funding or sponsorship was received for this study or publication of this article.

## Author contributions

C.S.P.B. and A.R. Conceptualization and designed the study. C.S.P.B., F.N. and A.R. Methodology. R.A., D.M., D.R., C.S.P.B and A.R. analyzed and interpreted the AST results. R.A., D.M., D.R., C.S.P.B. and A.R. Data curation. C.S.P.B. Formal analysis. C.S.P.B. drafted the original manuscript. F.N., and A.R. reviewed and edited the manuscript. F.N., and A.R. supervised the study. All authors reviewed and approved the final version of the manuscript.

## Conflict of interest

The authors declare no conflicts of interest.

